# Electrospun fibers for vaginal administration of tenofovir disoproxil fumarate and emtricitabine in the context of topical pre-exposure prophylaxis

**DOI:** 10.1101/2021.02.01.429162

**Authors:** Rute Nunes, Sarah Bogas, Maria João Faria, Hugo Gonçalves, Marlene Lúcio, Teresa Viseu, Bruno Sarmento, José das Neves

## Abstract

Women are particularly vulnerable to sexual HIV-1 transmission. Oral pre-exposure prophylaxis (PrEP) with tenofovir disoproxil fumarate and emtricitabine (TDF/FTC) is highly effective in avoiding new infections in men, but protection has only been shown moderate in women. Such differences have been associated, at least partially, to poor drug penetration of the lower female genital tract and the need for strict adherence to continuous daily oral intake of TDF/FTC. On-demand topical microbicide products could help circumventing these limitations. We developed electrospun fibers based on polycaprolactone (PCL fibers) or liposomes associated to poly(vinyl alcohol) (liposomes-in-PVA fibers) for the vaginal co-delivery of TDF and FTC, and assessed their pharmacokinetics in mice. PCL fibers and liposomes-in-PVA fibers were tested for morphological and physicochemical properties using scanning electron microscopy, differential scanning calorimetry and X-ray diffractometry. Fibers featured organoleptic and mechanical properties compatible with their suitable handling and vaginal administration. Fluorescent quenching of mucin *in vitro* – used as a proxy for mucoadhesion – was intense for PCL fibers, but mild for liposomes-in-PVA fibers. Both fibers were shown safe *in vitro* and able to rapidly release drug content (15-30 min) under sink conditions. Liposomes-in-PVA fibers allowed increasing genital drug concentrations after a single intravaginal administration when compared to continuous daily treatment with 25-times higher oral doses. For instance, the levels of tenofovir and FTC in vaginal lavage were around 4- and 29-fold higher, respectively. PCL fibers were also superior to oral treatment, although to a minor extent (approximately 2-fold higher drug concentrations in lavage). Vaginal tissue drug levels were generally low for all treatments, while systemic drug exposure was negligible in the case of fibers. These data suggest that proposed fibers may provide an interesting alternative or an ancillary option to oral PrEP in women.

## 1. Introduction

Global incidence of HIV infection remains at unacceptably high levels and is unlikely to significantly regress over the next decade [1]. Women in particular are largely vulnerable to sexual transmission due to various biological, behavioral and socio-demographic factors [2]. Protecting women from HIV/AIDS is, therefore, a health imperative. Oral pre-exposure prophylaxis (PrEP) emerged over the last decade as an effective approach to prevent the sexual transmission of HIV-1. Daily oral administration of antiretroviral drugs, namely the combination of tenofovir disoproxil fumarate/emtricitabine (TDF/FTC), has been shown highly efficacious in preventing HIV-1 transmission in men who have sex with men (MSM) [3]. However, outcomes of clinical trials involving heterosexual couples have been mixed regarding the efficacy of oral PrEP in women [4–7]. Adherence to a daily oral regimen has been pointed out as the main cause for failure, but other relevant factors may also be involved [8]. For instance, Cottrell *et al*. [9] showed that poor drug penetration of the lower female genital tract – particularly in the case of tenofovir (TFV) resulting from the hydrolysis of its prodrug TDF by plasma and tissue esterases – may limit the ability of oral PrEP to protect women. Furthermore, slow onset of protective drug levels and their rapid decline following discontinuation impair the implementation of more flexible regimens such as on-demand regimens that have been successfully introduced in MSM [10]. Typically, 6-7 consecutive daily doses of TDF/FTC are deemed necessary to achieve maximal protection in women [11]. Topical PrEP using vaginal microbicides could help tackling this limitation of oral TDF/FTC in achieving rapid protective drug levels at the genital tract mucosa and potentially allow on-demand use. In particular, the vaginal administration of TDF alone [12, 13] or in combination with FTC [14, 15] was able to effectively prevent infection in animal models.

Engineering suitable drug delivery platforms is a key step in the microbicide development process. Electrospun polymeric fibers have been proposed for vaginal drug delivery, particularly in the context of HIV-1 prevention, contraception and sexual health [16]. These systems are easy to produce at both the laboratory and industrial scales and are highly versatile. For instance, fibers can simultaneously incorporate different active compounds and be engineered to modulate drug release by using different materials or by changing their architecture [17–19]. Fiber mats can be used similarly to commercially available vaginal films, presenting advantages such as the possibility of self-administration without the need of an applicator, avoidance of extensive leakage typical of semisolid products, or minimal interference with sexual intercourse [20]. Preclinical data suggest that fibers allow achieving quick and extensive coverage of the vaginal epithelium, which could be crucial for providing effective protection from HIV-1 transmission [21]. Furthermore, an exploratory study conducted by performing focus group interviews found that fiber mats are likely to be well accepted by women for topical PrEP [22].

We hypothesize that vaginal TDF/FTC-loaded fibers may be interesting systems for on-demand topical PrEP. Two types of electrospun formulations, each loaded with both drugs and comprising either hydrophobic polycaprolactone (PCL) fibers or a composite of liposomes and hydrophilic poly(vinyl alcohol) (PVA) fibers, were developed. In the case of the composite, we attempted to explore liposomes not only for the purpose of retaining TDF and FTC in fibers, but also for taking advantage of the possible capacity of nanocarriers to improve microbicide drug distribution and retention in the vagina [23]. Both types of fibers were evaluated regarding the ability to provide potentially protective vaginal drug levels, namely by comparing single-dose pharmacokinetics (PK) of fibers to that of continuous use of oral PrEP with TDF/FTC in a murine model.

## 2. Materials and methods

### 2.1. Materials

TDF was purchased from Kemprotec (Cumbria, UK) and FTC from Sequoia Research Products (Pangbourne, UK). TFV was kindly supplied by Gilead Sciences (Foster City, CA, USA). PCL (80 kDa), sodium lauryl sulfate, purified type II mucin and thiazolyl blue tetrazolium bromide were acquired from Sigma-Aldrich Química (Sintra, Portugal), PVA (57-66 kDa) from Alfa Aesar (Kandel, Germany), and 1,2-dimyristoyl-*sn*-glycero-3-phosphocholine (DMPC), 1,2-dioleoyl-*sn*-glycero-3-phosphoethanolamine (DOPE) and cholesterol (CHL) from Avanti Polar Lipids (INstruchemie, Delfzyl, The Netherlands). Truvada^®^ (Gilead Sciences, Lisbon, Portugal) was acquired from a local pharmacy. All other reagents and solvents were of analytical grade or equivalent.

### 2.2. Preparation and characterization of liposomes

Liposomes were prepared using the thin film hydration technique by modifying a previously reported method [24]. Briefly, DMPC, DOPE and CHL at a molar ratio of 7:2:1 were dissolved in chloroform and the resulting solution was evaporated at 37 °C under a constant stream of nitrogen. The obtained film was hydrated with ultrapure water above the lipid main phase transition temperature and containing 40 μM of TDF and 28 μM of FTC. The lipid suspension was vortexed to produce liposomes with a final concentration of 4 mM, which were then sonicated in an ultrasonic bath for one minute. Liposomes were characterized for mean hydrodynamic diameter and zeta potential at 25 °C using a Zetasizer Nano ZS (Malvern Panalytical, Malvern, UK).

### 2.3. Production of fibers

PCL-based fibers containing TDF/FTC (further referred to as ‘PCL fibers’) and PVA-based fibers incorporating TDF/FTC-loaded liposomes (further referred to as ‘liposomes-in-PVA fibers’) were produced by electrospinning. Both polymers were dissolved in suitable solvent systems to a final concentration of 7-10% (*w/v). A* mixture of chloroform and methanol (3:2 in volume) at room temperature was used for PCL, while PVA was dissolved in an aqueous dispersion of liposomes at 80 °C. TDF and FTC (or drug-loaded liposomes) were then added to a final concentration of 4.0% (*w/w*) and 2.8% (*w/w*), respectively. An in-house electrospinning setup composed by a CZE2000 high voltage power supply (Spellman, Hauppauge, NY, USA), a 5 mL plastic syringe coupled to a blunt-end 22G (PCL fibers) or 19G (PVA fibers) needle, a single syringe pump (KD Scientific, Holliston, MA, USA), and a grounded drum collector rotating at 100 rpm was used to produce fibers. Precursor solutions were pumped at 0.6 mL·h^−1^ (PCL) or 0.1 mL·h^−1^ (PVA) in the presence of a 15 kV (PCL fibers) or 20 kV (PVA fibers) electrical field and collected at a distance of 11 cm. Resulting fiber mats were dried and maintained in a vacuum desiccator until used. PCL fibers and liposomes-in-PVA fibers without drugs, as well as PVA fibers without liposomes and drugs, were similarly obtained by omitting drug and/or liposome incorporation during production.

### 2.4. Physicochemical, mechanical and biopharmaceutical characterization of fibers

The morphology of fibers was analyzed by scanning electron microscopy (SEM) using a Nova NanoSEM 200 (FEI, Hillsboro, OR, USA) at an acceleration voltage of 15 kV. Cryogenic fractured samples were sputter coated with gold before analysis. Cross-section diameter values of fibers were determined from acquired images using the Digimizer^®^ software (MedCalc Software, Ostend, Belgium). A minimum of 100 measurements were performed for each fiber type. The apparent porosity percentage (*P*%) of fiber mats was estimated from thickness measurements according to the following equation:

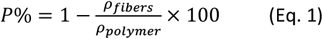

where *ρ_polymer_* is the density of PCL (1.15 g·cm^−3^) or PVA (1.26 g·cm^−3^) and *ρ_fibers_* is the apparent density of fiber mats obtained by dividing the mass of a sample by its volume.

Thermal transitions and thermodynamic parameters of the fibers were analyzed by differential scanning calorimetry (DSC). Measurements were carried out under nitrogen atmosphere using a DSC 214 Polyma (NETZSCH, Selb, Germany), as previously described [25]. Thermograms were obtained at a heating rate of 10 °C·min^−1^ in the range of 0-100 °C for PCL fibers and 150-250 °C for liposomes-in-PVA fibers. Thermodynamic parameters were calculated using the Proteus^®^ 7.1 software (NETZSCH).

X-ray diffraction patterns were acquired with the micro-focus wide-angle X-ray scattering beam line (MiNaXS) of PETRA III (DESY, Hamburg, Germany) using monochromatic synchrotron X-ray radiation (energy of approximately 15 keV and wavelength in the range of 0.82 Å). To observe the complete scattering pattern, a 2D pixel detector PILATUS 1M (Dectris, Baden-Daettwil, Switzerland) was used, with a pixel size of (172 × 172) μm^2^.

The mechanical properties of fiber mats were studied using an AG-IS universal testing machine (Shimadzu, Kyoto, Japan). Stress-strain tests were carried out with a 10 N load cell and a deformation speed of 10 mm·min^−1^ until rupture of fiber mat samples (10 mm in width and 25 mm in length). Young’s modulus (*E*) was estimated by linear fitting of stress-strain profiles for 0-7% of maximum stress.

Mucoadhesive potential of fibers was assessed by estimating their quenching effect on the intrinsic fluorescence of mucin [24]. Fiber mat samples were immersed in an aqueous mucin dispersion (0.2 mg·mL^−1^) for 180 min at 37 °C and 100 rpm. Fluorescence measurements of the mucin suspension before and after contact with fibers were carried out at excitation/emission wavelengths of 270 nm/300-500 nm using a LS 50B fluorimeter (Perkin Elmer, Waltham, MA, USA).

*In vitro* release of TDF and FTC from fibers was assessed in lactate buffer (pH 4.5) containing 12 mM SLS (micellar medium) in order to allow establishing sink conditions while mimicking the amphiphilic content of the cervicovaginal fluids [26]. Briefly, fiber samples (3 × 2 cm) were immersed in 30 mL of medium and maintained at 37 °C under shaking conditions (120 rpm). Medium samples (1 mL) were collected at pre-determined time points and replaced with fresh medium, before being assayed by a UV/Vis spectrophotometric derivative method in the range of 200-400 nm [27]. *In vitro* drug release profiles were fitted to commonly used kinetics models (first order, Korsmeyer-Peppas and Gallagher-Corrigan) using Prism^®^ 5 software (GraphPad Software, La Jolla, CA, USA).

Cytotoxicity of fibers to two female genital cell lines was assessed using the MTT metabolic activity assay, as previously described [28]. Ca Ski cervical and HEC-1-A endometrial cells (ATCC, Manassas, VA, USA), were maintained in RPMI 1640 medium or McCoy’s 5A modified medium, respectively, supplemented in both cases with 10% (*v/v*) fetal bovine serum (Biochrom GmbH, Berlin, Germany), 100 U·mL^−1^ penicillin and 100 μg·mL^−1^ streptomycin (BioWest, Nuaillé, France) under standard conditions (37 °C, 5% CO2 and 95% RH). Extracts were prepared by immersing fiber samples in culture medium at a mat surface area-to-volume of medium of 1 cm^2^·mL^−1^ during 24 h at 37 °C. Extracts were added to cells (10^4^ per well pre-incubated for 24 h in 96-well plates) and allowed to incubate for 24 h. Cells were then washed twice with phosphate buffered saline (pH 7.4) before adding MTT at 0.5 mg·mL^−1^ in medium and incubating for 4 h under standard conditions. Formazan crystals were dissolved with DMSO and cell viability was determined by measuring the absorbance at 570 nm.

Extracts of 1% (*w/v*) Triton^®^ X-100 and plain media were used as controls. Also, extracts of a commercial contraceptive film containing 28% nonoxynol-9 (VCF^®^ Vaginal Contraceptive Film, Apothecus, Oyster Bay, NY, USA) were tested for comparison purposes.

### 2.5. Pharmacokinetics of TDF/FTC-loaded fibers

The concentrations of TDF, TFV and FTC in vaginal lavage and tissue, as well as blood plasma, were determined between 15 min and 24 h after either intravaginal administration of fibers containing TDF/FTC or oral administration of the drug combinations to 8-12 weeks old female ICR mice. All procedures and experiments were performed using mice bred at the i3S animal facility, approved by the Ethics Committee at IUCS-CESPU (process no. 01/ORBEACESPU/2014) and conducted under the European Directive 2010/63/EU guidance, as previously detailed [29, 30]. Mice were pre-treated subcutaneously with 3 mg of medroxyprogesterone (Depo-Provera^®^, Pfizer, Porto Salvo, Portugal) at seven and three days before administration of TDF/FTC in order to induce a diestrus-like state. Square-shaped fiber mats with approximately 20-25 mm^2^ and containing of 0.07 mg of TDF and 0.05 mg of FTC were folded in half and administered intravaginally to conscientious animals with the aid of tweezers [30]. Oral gavage of TDF/FTC to conscientious mice was conducted after pulverization and dispersion of Truvada^®^ tablets in water. Five daily administrations of TDF (62 mg·kg^−1^) and FTC (41 mg·kg^−1^) were performed. Animals were allowed to move freely throughout all experiments, with free access to water and food.

After a single intravaginal administration of fibers or following the fifth oral administration of TDF/FTC, mice were euthanized at pre-determined time points (15 min, 1 h, 4 h and 24 h) by inhalational isoflurane overdose, followed by intracardiac exsanguination. Blood was collected into Vacuette^®^ tubes with K3EDTA (Greiner Bio-One, Kremsmünter, Austria) and plasma was recovered by centrifugation (1,180 ×*g*, 10 min) at 4 °C. The vagina was washed four times with 50 μL of normal saline using a micropipette, and the full recovered fluid centrifuged (13,414 ×*g*, 10 min) at 4 °C before collection of supernatant (lavage). Upon necropsy, the vagina was dissected and cut open along its longest axis. Visible residues of fibers were removed in the case of animals treated intravaginally before tissue collection. All samples (plasma, lavage and tissue) were stored at −80 °C until further processing. Drug assay was conducted by liquid chromatography-tandem mass spectrometry (LC-MS/MS), as detailed in *Supporting Information (S1. Supporting Methods*). Data were used to determine: (i) the maximum observed concentration (*C*_max_) and time at which it occurred (*t*_max_); (ii) the area-under-the-curve between 15 min and 24 h (AUC_0.25-24h_), as calculated by the trapezoidal rule using Prism^®^ 5 software; and (iii) the relative bioavailability (*F*_rel_), defined as the ratio between AUC_0.25-24h_ values obtained for fibers and oral TDF/FTC, without dose adjustment. Groups of five animals were used per treatment and time point.

### 2.6. Statistical analysis

Experiments were performed in triplicate unless otherwise noted. Data are presented as mean ± standard deviation (SD), except in the case of PK results for which error is expressed as the standard error of the mean (s.e.m.). Multiple group comparisons were performed by one-way ANOVA with Tukey’s HSD post-hoc using Prism^®^ 5 software. Values of *p* < 0.05 were assumed as denoting significant differences.

## 3. Results and discussion

### 3.1. Physicochemical and mechanical characterization of TDF/FTC-loaded fibers

Both types of TDF/FTC-loaded fibers were successfully produced by electrospinning. Drug-loaded liposomes featuring diameter of 211 ± 24 nm and zeta potential of −0.67 ± 0.01 mV were incorporated into PVA fibers. Drug content of liposomes-in-PVA fibers was 0.28 mg·cm^−2^ in TDF and 0.20 mg·cm^−2^ in FTC, while PCL fibers contained 0.34 mg·cm^−2^ of TDF and 0.24 mg·cm^−2^ of FTC. PCL and liposomes-in-PVA fiber meshes were glossy or pale white, respectively, homogeneous, and soft to the touch (*Supporting Information, Figure S1*). Thickness of PCL and liposomes-in-PVA fiber mats was 364 ± 2.74 μm and 133 ± 4.67 μm, respectively, while *P*% was similar in both cases and estimated at 88-90% (*Supporting Information, Figure S1*). SEM imaging revealed that both types of fibers were continuous and had a smooth surface (**Figure 1A**). Values for cross-section diameter of PCL fibers and liposomes-in-PVA fibers containing TDF and FTC were 726 ± 279 nm and 222 ± 86 nm, respectively. PCL fibers were not only thicker, but also more heterogeneous (**Figure 1B**). The incorporation of drugs and drug-loaded liposomes was shown to substantially modify the appearance of both types of fibers. PCL fibers without TDF and FTC were roughly 20% thinner (591 ± 191 nm) than drug-loaded counterparts (*Supporting Information, Figure S2*). PCL (logP = 4.03) is highly lipophilic and unable to dissolve moderately lipophilic/hydrophilic drugs, such as TDF (logP = 1.25) and FTC (logP = −0.43). This may dictate the disruption of PCL crystalline domains with consequent fiber enlargement. Conversely, PVA without TDF/FTC-loaded liposomes featured larger a cross-section diameter value of 460 ± 118 nm (*Supporting Information, Figure S2*). PVA (logP = 0.26) is not only able to molecularly disperse TDF and FTC, but the incorporation of liposomes may also have led to better polymeric alignment and consequent decrease in fiber diameter. Interestingly, drug-loaded liposomes-in-PVA fibers featured distinctive points of enlargement (roundish bulges), possibly due to the presence of agglomerated liposomes (**Figure 1A**). This effect has been previously reported by others [31] and was particularly noticeable in the case of liposomes-in-PVA fibers without drugs (*Supporting Information, Figure S2*). The reduced amount of enlarged points in drug-loaded liposomes-in-PVA fibers may be related to the ability of the active compounds (especially FTC) to disturb lipid packing [24], thus making liposomes possibly more flexible and featuring less roundish bulges.

**Figure 1.**
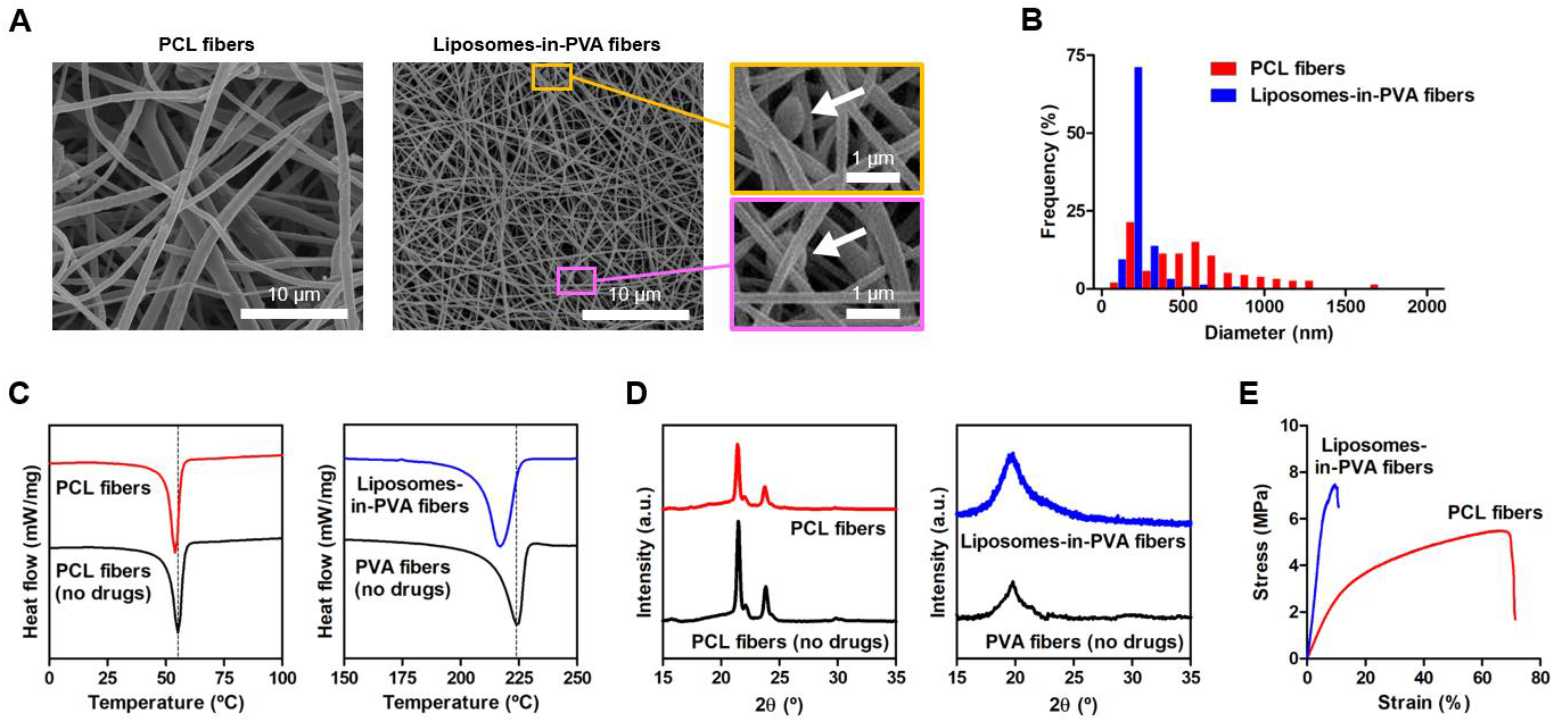
Physicochemical and mechanical properties of TDF/FTC-loaded fibers. (A) Representative SEM images (insets highlight discrete points of enlargement along liposomes-in-PVA fibers). (B) Cross-section diameter distribution. (C) DSC thermograms and (D) X-ray diffractograms of fibers (with or without the incorporation of drugs/drug-loaded liposomes). (E) Representative stress-strain curves.

DSC analysis was conducted in order to infer about the compatibility of fiber ingredients and to analyze the effect of drugs/liposomes on fiber structure. Thermograms underscore the dominance of PCL or PVA, namely by featuring single endothermal peaks resulting from polymer fusion (**Figure 1C**). Melting temperature (*T*_m_) values and correspondent enthalpy change (*ΔH*) are presented in **Table 1** and overlap typical values for PCL or PVA [32, 33]. Still, mild reduction of *T*_m_ and Δ*H* were observed for PCL-based fibers upon drug incorporation, corresponding to a 5% loss in crystallinity. This suggests lesser fiber stiffness when drugs are present, which is consistent with the different lipophilicity of drugs and polymer. Thermograms of liposomes-in-PVA fibers did not feature typical melting endotherms of drugs (TDF at 115 °C [34] and FTC 180 °C [35]) or lipids, thus suggesting good miscibility/incorporation of drugs and liposomes in the polymeric matrix. However, a small reduction in *T*_m_ was observed upon drug/liposome addition, which was accompanied by a marked increase in the crystallinity of the polymeric matrix. This appears to indicate that liposomes favored the homogeneous distribution of drugs in PVA fibers, as well as their compatibility.

**Table 1.** Thermodynamic parameters and mechanical properties of fibers, as evaluated by DSC and tensile stress tests. Results for mechanical properties are presented as mean ± SD (*n* = 3).

| Fibers | Thermodynamic parameters |  |  | Mechanical properties |  |  |
| --- | --- | --- | --- | --- | --- | --- |
| | $T_m$ ($^\circ\text{C}$) | $\Delta H$ ($\text{J}\cdot\text{g}^{-1}$) | $\Delta C_{\text{crys}}$ (%) | $TS_{\text{max}}$ (MPa) | $\epsilon$ (%) | $E$ (MPa) |
| PCL fibers (no drugs) | 55.2 | 51.3 | – | $12.8 \pm 0.8$ | $99.8 \pm 9.2$ | $41.3 \pm 0.4$ |
| PCL fibers | 54.0 | 48.5 | -5 | $5.5 \pm 0.1$ | $67.8 \pm 9.0$ | $28.9 \pm 0.3$ |
| PVA fibers (no drugs) | 225.0 | 50.5 | – | $9.5 \pm 1.5$ | $11.2 \pm 1.2$ | $64.2 \pm 0.6$ |
| Liposomes-in-PVA fibers | 216.9 | 75.2 | +49 | $7.3 \pm 1.2$ | $10.3 \pm 1.5$ | $86.6 \pm 0.8$ |

$T_m$ – melting point temperature; $\Delta H$ – enthalpic variation; $\Delta C_{\text{crys}}$ – variation of crystalline structure (calculated as $\Delta C_{\text{crys}} = 100 \times \Delta H_{\text{fibers with drugs}} / \Delta H_{\text{fibers without drugs}}$ ); $TS_{\text{max}}$ – Maximum tensile strength at break; $\epsilon$ – elongation at break; $E$ – Young's modulus.

We next tested fibers by X-ray diffraction in order to confirm crystallinity changes inferred from DSC data. Diffractograms are shown in **Figure 1D**. PCL fibers featured two diffraction peaks at 2θ = 21.4 ° and 2θ = 23.8 ° corresponding to the reflections according to the planes (110) and (200), respectively [36, 37]. Both are indicative of the distinctive orthorhombic crystal lattice of the polymer, being only slightly decreased when TDF/FTC were incorporated. This mild loss of the crystallinity of PCL is in agreement with DSC results. In the case of PVA-based fibers, diffractograms were dominated by a characteristic peak at 2θ = 19.3 ° [38]. The higher intensity of this peak after the incorporation of TDF/FTC-loaded liposomes supports an increase in crystallinity of fibers and confirms DSC results.

Changes in crystallinity due the incorporation of drugs/liposomes are deemed acceptable as long as they do not compromise the mechanical resistance and structural integrity of mats. Therefore, we performed uniaxial tensile tests (**Figure 1E**) and calculated parameters such as elongation (*ε*) and tensile strength (*TS*_max_) at break, as well as elastic modulus (Young’s modulus; *E*), in order to evaluate the mechanical properties of fibers. Considerable differences were observed when drugs/liposomes were present or not (**Table 1** and *Supporting Information, Figure S3*). Variations in crystallinity were positively correlated with values of *E* for both types of fibers. PCL fiber mats containing drugs demonstrated a reasonable balance between flexibility and hardness, being able to withstand over 50% extensional strain. PVA-based fiber mats were less flexible (lower *ε* values) than PCL fibers, but the incorporation of TDF/FTC-loaded liposomes did not affect this feature. From a practical point of view, both types of fiber mats could be considered as resistant to relevant handling (*e.g*., occurring during manufacturing, transport or administration), namely when compared to vaginal films designed for use as microbicides (*E* = 5.4-7.8 MPa [39]).

### 3.2. Biopharmaceutical properties of TDF/FTC-loaded fibers

The ability of a drug delivery system to associate with mucin at the nanoscale provides a valuable indicator of its putative mucoadhesive properties [40]. We assessed the capacity of fibers to quench the intrinsic fluorescence of mucin by establishing hydrophobic interactions with tryptophan residues. PCL-based fibers were able to reduce fluorescence intensity (**Figure 2A**), thus suggesting strong binding to mucin. The quenching effect was particularly pronounced for fibers containing TDF/FTC, which was related to the ability of drugs themselves to interact with tryptophan (*Supporting Information, Figure S4*). Additional SEM imaging indicated that mucin was able to intimately coat PCL-based fibers, while X-ray diffraction experiments showed decreased intensity in typical peaks of this type of fibers when mucin was present (*Supporting Information, Figure S4*). Altogether, these results suggest that PCL fibers are highly mucoadhesive [41]. Conversely, liposomes-in-PVA fibers did not appear to interact with mucin, except when TDF/FTC were included (**Figure 2A**). This seems to indicate that PVA-based fibers feature low mucoadhesiveness. Fluorescence quenching experiments were again reinforced by results of X-ray diffraction patterns for PVA-based fibers in the presence or absence of mucin (*Supporting Information, Figure S4*). Overall, and assuming the quenching percentage as a quantitative indicator of mucoadhesiveness, PCL fibers were approximately 10-fold more mucoadhesive than liposomes-in-PVA fibers (*Supporting Information, Figure S5*). These results, however, should be interpreted with caution. Quenching of the intrinsic fluorescence of mucin mostly assesses hydrophobic interactions [41], which have been estimated as quite relevant in defining the mucoadhesiveness of vaginal formulations [42]. Nevertheless, other types of interactions (*e.g*., electrostatic and hydrogen bonding) that are important in defining the adhesiveness of hydrophilic polymers may have been underestimated.

**Figure 2.**
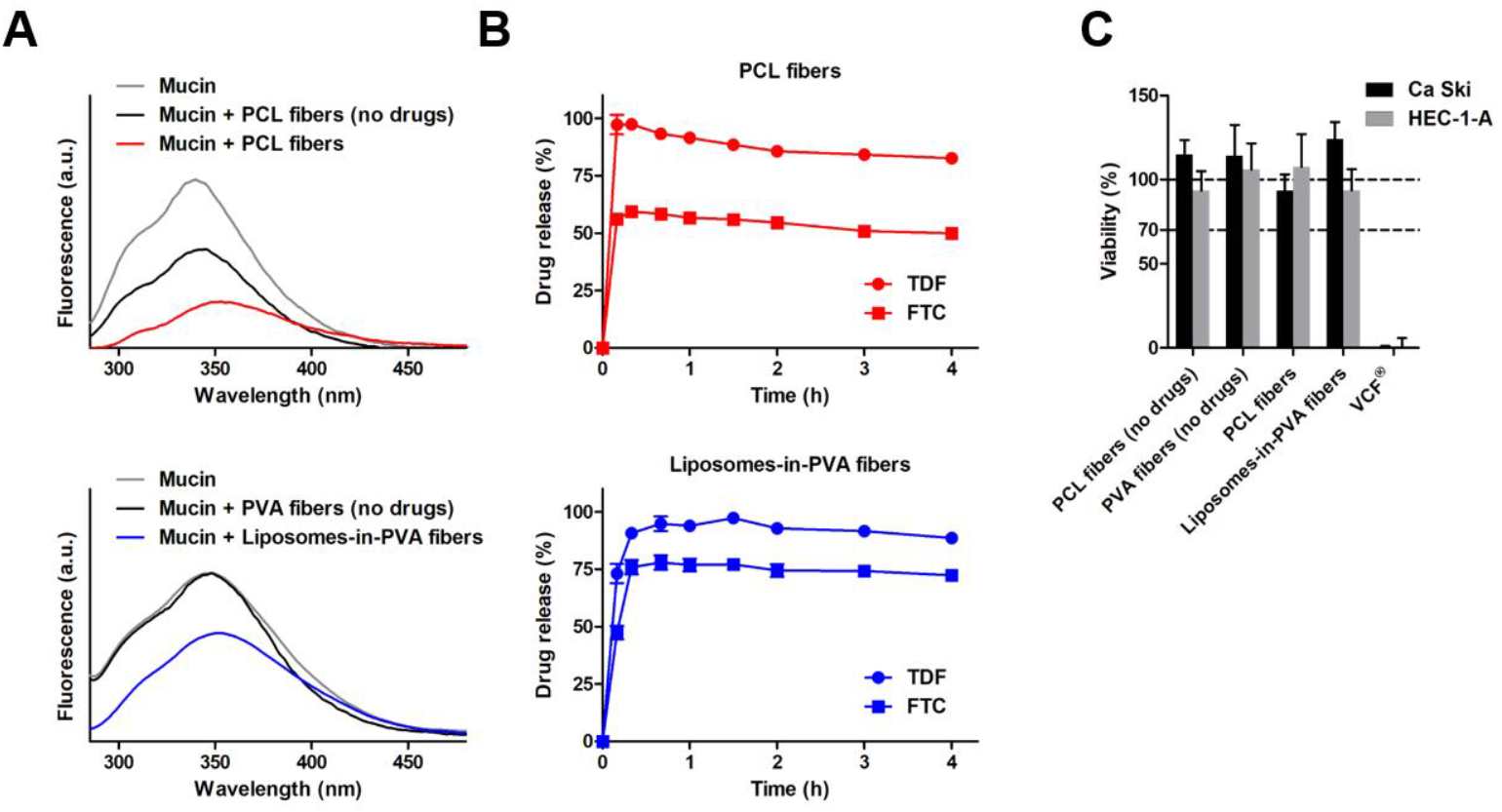
Biopharmaceutical properties of TDF/FTC-loaded fibers. (A) Fluorescence emission spectra of mucin before and after incubation with fibers. (B) Drug release profiles of drugs from fibers in micellar medium (pH 4.5) at 37 °C. (C) Viability of Ca Ski and HEC-1-A cells upon incubation with fibers (with or without the incorporation of drugs/drug-loaded liposomes). Results in (B) and (C) are presented as mean ±SD (*n* = 3).

Next, we evaluated the ability of fibers to release TDF and FTC in micellar medium at pH 4.5. Drug release from PCL fibers was fast (within 15-30 min), even if roughly half the amount of FTC was retained up to at least 4 h (**Figure 2B**). While the rapid and complete release of TDF suggests that this drug is located mostly at the surface of the system, the profile of FTC points to a more homogeneous distribution within the PCL matrix [43]. Previous studies showed that TDF typically presents burst release from electrospun fibers composed of hydrophobic polymers such as (poly(lactic-*co*-glycolic acid) (PLGA), poly(lactide-*co*-caprolactone), or a 20:80 blend of PCL and PLGA [19, 43]. Noticeably, the apparent decrease in released fractions of TDF from PCL fibers up to 4 h were likely related with the hydrolysis of the two isopropyloxycarbonyloxymethyl groups of this prodrug in aqueous media, as previously described by Carson *et al*. [19] Liposomes-in-PVA fibers presented similar release profiles to PCL fibers, although the amount of retained FTC was lower (approximately 25%). This fraction of FTC is likely intimately associated with liposomes as PVA-based fibers are highly soluble and prone to rapidly release payloads. Release profiles of both types of fibers were best fitted to the Gallagher-Corrigan model (*R*^2^_adjusted_ ≥ 0.979; *Supporting Information, Tables S1 and S2*), which is indicative of typical biphasic behavior (initial burst followed by slow release) of biodegradable polymeric system [44]. Overall, rapid drug release observed for PCL fibers and liposomes-in-PVA fibers may be considered as beneficial for developing on-demand microbicides, since pericoital administration requires that protective levels of TDF and FTC are achieved in a very short amount of time. Although not explored, fibers developed in this work may still be useful for developing sustained release products, namely by stacking drug-loaded and non-medicated sheets of fiber, as previously proposed by other researchers [45, 46].

Lastly, we assessed the toxicity potential of fibers using well-established *in vitro* genital cell models. Extracts of fibers were shown not to be cytotoxic (**Figure 2C**). Cell viability was maintained above 90% for PCL fibers and liposomes-in-PVA fibers – well above the 70% threshold that is usually considered when testing biomedical materials and devices [47] -,contrasting with the results for VCF^®^ film extracts. These data are in agreement with the recognized safety of PCL and PVA, as well as the relatively low toxicity potential of TDF and FTC. For instance, the maximum concentrations achievable for TDF and FTC in fiber extracts were 140 μg·mL^−1^ and 100 μg·mL^−1^, respectively, which are below the half-maximal cytotoxic concentration (CC50) values determined for these drugs in the HeLa cervical cell line (231 μg·mL^−1^ and > 300 μg·mL^−1^, respectively) [28].

### 3.3. Pharmacokinetics of TDF/FTC-loaded fibers

We conducted *in vivo* experiments in a non-infectious animal model in order to compare local and systemic PK of intravaginally administered fibers with oral TDF/FTC. The vaginal doses of TDF (0.07 mg) and FTC (0.05 mg) were selected in accordance with previous studies demonstrating efficacy in preventing vaginal HIV-1 transmission in humanized mice [12, 15]. Truvada^®^ was used in order to better mimic real-world usage of TDF/FTC. Tablets can be crushed and dispersed in water immediately before intake without affecting PK according to information included in the ‘Summary of Product Characteristics’. TDF/FTC was administered by gavage at 61.5 mg of TDF and 41 mg of FTC per kg of body weight (animals weighing about 30 g), which correspond to roughly 25-times the doses administered intravaginally. These were selected by considering the human doses for oral PrEP and back-calculating the amount of each drug for mice using allometric scaling, as defined by the FDA [48]. While the PK of fibers were assessed after a single dose, five daily doses were administered to mice in order to simulate continuous oral PrEP and achieve steady-state drug levels [49].

Drug concentrations – including those of TFV resulting from the hydrolysis of TDF – were determined in vaginal lavage and tissue, as well as in blood plasma, and are presented in **Figure 3**. Sampling times were 15 min, 1 h, 4 h and 24 h post-administration, which allow assessing representative short-term, medium-term, long-term and terminal levels of topical microbicides in mice [29, 30]. Results for vaginal lavage (**Figure 3A**) showed that concentrations of TDF, TFV and FTC were generally higher for liposomes-in-PVA fibers as compared to oral PrEP, particularly at earlier time points, which suggests that these fibers could rapidly provide protective levels upon a single administration. Differences between liposomes-in-PVA fibers and oral TDF/FTC can even be better assessed by comparing values of AUC_0.25-24h_ (**Table 2**). For instance, TFV and FTC levels were roughly 4- and 29-times higher, respectively. Differences were also significant for TDF, for which no detectable levels were registered for oral administration. PCL fibers also allowed achieving higher drug concentrations than oral TDF/FTC, although differences were less pronounced. This could be attributed, at least in part, to the inability of hydrophobic fibers to fully release drugs in the reduced amount of fluids present in the vagina of mice. Indeed, 4 and 3 animals (out of 5) still presented readily identifiable PCL fiber mats at 15 min and 1 h post-administration, respectively (*Supporting Information, Tables S3*). The capacity of both fibers to allow achieving tangible vaginal levels of TDF (contrasting with non-detectable levels for oral treatment) are particularly relevant due to the enhanced capacity of the prodrug to yield intracellular levels of the active metabolite TFV diphosphate as compared to TFV [50]. Overall, the results for vaginal lavage seem to support that the administration of TDF/FTC-loaded fibers may be able to provide immediate protective concentrations of both drugs that are superior to oral TDF/FTC.

**Figure 3.**
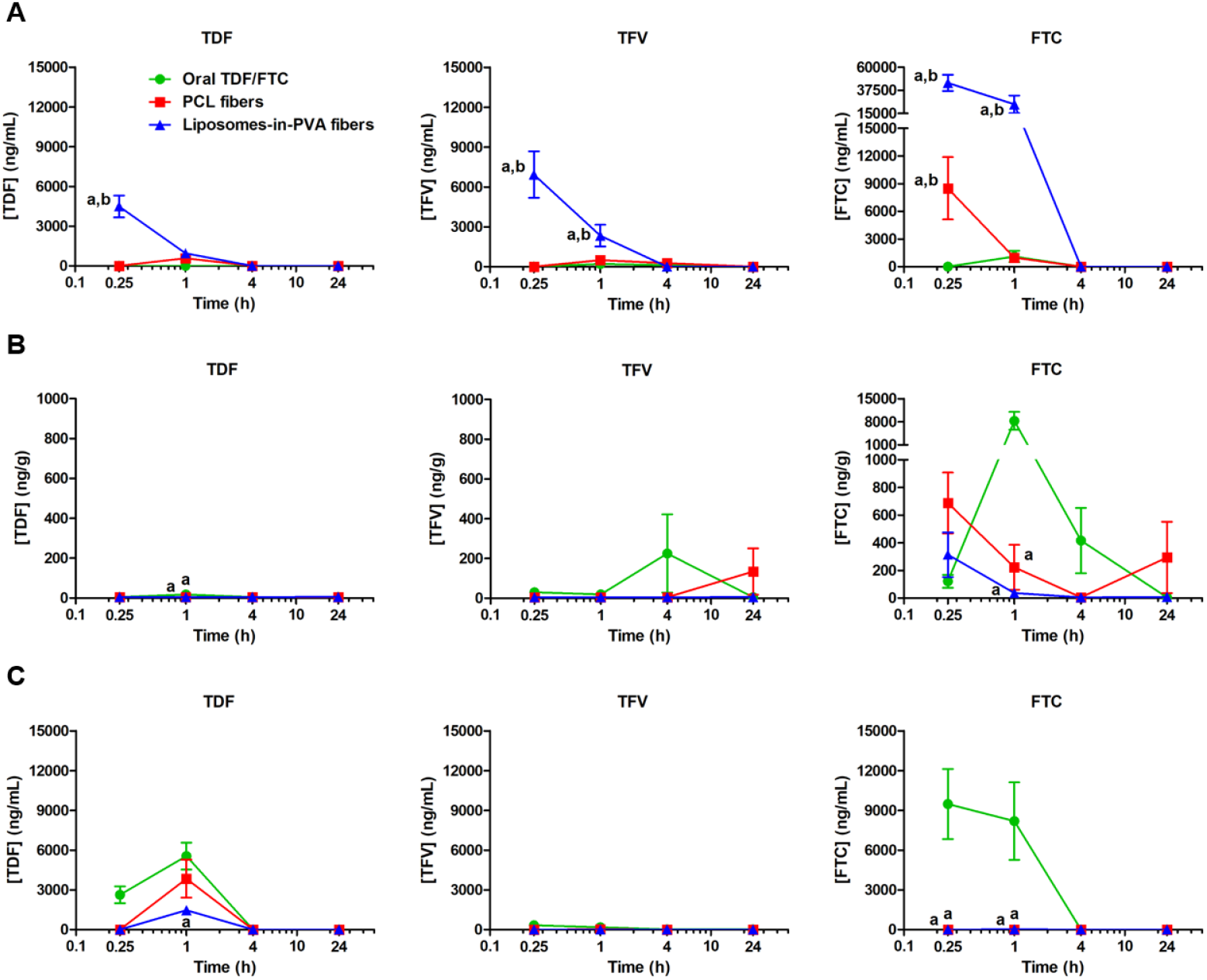
PK profiles of fibers and oral TDF/FTC. Time-dependent concentrations of TDF, TFV and FTC in different biological samples upon administration of vaginal fibers or oral TDF/FTC. Concentrations of drugs in (A) vaginal lavage, (B) vaginal tissue and (C) blood plasma are presented as mean ± s.e.m. (*n* = 5). ^(a)^ and ^(b)^ denote statistically significant differences (*p* < 0.05) when comparing with oral TDF/FTC or the other type of fibers, respectively.

**Table 2.** PK parameters for TDF, TFV and TDF in vaginal lavage (VL), vaginal tissue (VT) and blood plasma (BP) after vaginal administration of drug-loaded fibers or oral administration of TDF/FTC. Results for *C*_max_ and *t*_max_ are presented as mean ± s.e.m. (*n* = 5).

| Drugs | Formulation | $C_{\max}$ (ng·mL<sup>-1</sup>) <sup>(a)</sup> | | | $t_{\max}$ (h) | | | AUC<sub>0.25-24h</sub> (ng·h·mL<sup>-1</sup>) <sup>(b)</sup> | | | $F_{\text{rel}}$ | | |
| --- | --- | --- | --- | --- | --- | --- | --- | --- | --- | --- | --- | --- | --- |
|  |  | VL | VT | BP | VL | VT | BP | VL | VT | BP | VL | VT | BP |
| TDF | Oral TDF/FTC | ND | 17.9 ± 3.6 | 5,550 ± 1,013 | ND | 1 | 1 | 0 | 142.9 | 11,392 | – | – | – |
| | Vaginal PCL fibers | 584 ± 103 | ND | 3,848 ± 1,435 | 1 | ND | 1 | 1,096 | ND | 7,216 | ND | ≈0 | 0.6 |
| | Vaginal liposomes-in-PVA fibers | 4,488 ± 820 | 6.2 ± 0.7 | 1,474 ± 230 | 0.25 | 24 | 1 | 3,470 | 130.7 | 2,763 | ND | 0.9 | 0.2 |
| TFV | Oral TDF/FTC | 205 ± 72 | 225 ± 197 | 326 ± 162 | 1 | 4 | 0.25 | 1,763 | 2,690 | 818 | – | – | – |
| | Vaginal PCL fibers | 492 ± 64 | 135 ± 116 | 14.8 ± 4.0 | 1 | 24 | 1 | 3,981 | 1,415 | 26.8 | 2.3 | 0.5 | 0.03 |
| | Vaginal liposomes-in-PVA fibers | 6,929 ± 1,746 | 6.2 ± 0.7 | 7.4 ± 3.1 | 0.25 | 24 | 1 | 6,979 | 130.7 | 13.9 | 4.0 | 0.05 | 0.02 |
| FTC | Oral TDF/FTC | 1,115 ± 618 | 8,301 ± 236 | 9,475 ± 2,646 | 1 | 4 | 0.25 | 2,091 | 20,447 | 18,917 | – | – | – |
| | Vaginal PCL fibers | 8,498 ± 3,377 | 688 ± 220 | ND | 0.25 | 0.25 | ND | 5,040 | 3,670 | ND | 2.4 | 0.2 | ≈0 |
| | Vaginal liposomes-in-PVA fibers | 44,793 ± 7,944 | 314 ± 145 | 15.5 ± 15.5 | 0.25 | 0.25 | 1 | 61,564 | 325 | 29.1 | 29.4 | 0.02 | 0.002 |

<sup>(a)</sup> ng·g<sup>-1</sup> for VT; <sup>(b)</sup> ng·h·g<sup>-1</sup> for vaginal tissue. ND – not determined.

While results for lavage were quite distinct, data analysis for vaginal tissue was hindered by considerable interindividual variability and generally low drug concentrations in samples (**Figure 3B**). There was, however, a trend for higher drug levels in the case of oral TDF/FTC, which were significant at 1 h post-administration in the case of TDF and FTC. This suggests some degree of drug accumulation in tissue resulting from continuous oral use or simply poor mucosal permeation of intravaginally administered drugs. TDF, TFV and FTC are BCS class III compounds and substrates of important efflux transporters such as P-glycoprotein (TDF and TFV), Mrp1 (FTC) or Mrp4 (TFV), which are highly expressed in the cervicovaginal mucosa of mice [51–53]. PK parameters further reinforce the reduced ability of fibers to yield lower drug concentrations in tissue (**Table 2**). This was particularly apparent for FTC (*F*_rel_ = 0.02-0.2), while in the case of TFV or its prodrug, fibers could achieve similar levels of one of these compounds (*F*_rel_ (TDF) = 0.9 for liposomes-in-PVA fibers and *F*_rel_ (TFV) = 0.5 for PCL fibers). Noteworthy, TFV levels in vaginal tissue following oral administration were in general agreement with results reported for TDF administered orally to female BALB/c mice (AUC_2-24h_ = 9,681 ng·h·g^−1^ and *t*_max_ between 2 h and 8 h) [49], with variations likely reflecting differences in sample collection time points. In the case of vaginal administration of TDF and FTC, no comparable regional PK data have been previously described.

Data for blood plasma indicate low systemic drug exposure with the use of topical fibers, as denoted by their values of AUC_0.25-24h_ and *F*_rel_ (**Table 3**). It is worthwhile mentioning that one of the defining advantages of microbicides over oral PrEP relates with the principle of blocking transmission at the mucosal site and abbreviate the onset of possible adverse effects related with high systemic exposure to active payloads [54]. The blood plasma profile obtained for TFV with oral TDF/FTC was consistent with data previously found for the oral administration of TDF to female BALB/c mice, namely concerning the early onset of *C*_max_ [49, 55]. However, AUC values for similar oral doses were widely variable among studies: Veselinovic *et al*. [49] reported AUC_2-48h_ = 17,278 ng·h·mL^−1^ (12,251 ng·h·mL^−1^ when tested in humanized mice), while Ng *et al*. [55] reported AUC_0.5-24h_ = 3,984 ng·h·mL^−1^. Again, these variations probably reflect methodological differences between studies.

Apart from the discussion regarding relative drug levels in different biological samples for considered administration routes and/or fibers, it seems important to infer on how absolute values could translate into protection from HIV-1 transmission. Correlation between regional PK and efficacy for topical TDF/FTC has not yet been fully established. However, data from Gallay *et al*. [15] suggest that vaginal lavage concentrations of at least 4 ng/mL for FTC and 500 ng/mL for TDF (assuming a washing protocol similar to the one used in this work) are associated with full prevention of viral transmission in humanized mice. In our study, mean levels of FTC in lavage between 15 min and 1 h post-administration were at least around 6,000- and 250-times higher than the previous protective threshold in the case of liposomes-in-PVA fibers and PCL fibers, respectively. As for oral treatment, the concentration of FTC in lavage at 1 h was considerably above 4 ng/mL, but could not be guaranteed early on following administration. Mean levels of TDF in lavage were also maintained at least 2-times above the threshold reported by Gallay *et al*. for liposomes-in-PVA fibers between 15 min and 1 h. This was not the case for PCL fibers at 15 min. TDF in lavage was undetectable in the case of oral administration from 15 min to 24 h, which could reflect the conversion of the prodrug into TFV by esterases following oral absorption [56]. Furthermore, measurable plasmatic levels of TDF and TFV suggest that these drugs actually have poor ability to concentrate in the vaginal tissue of mice, thus paralleling what has been reported for women [9].

This study presents a few limitations that need to be considered when interpreting its findings. Firstly, PK data discussed in this manuscript reflect the analysis of a limited amount of time points that, nonetheless, typically reflect the dynamics of vaginal microbicide drug delivery in mice, as discussed above. Secondly, we did not evaluate tissue levels of active metabolites of TDF/TFV and FTC, namely TFV diphosphate and FCT triphosphate. However, preclinical (from mice) and clinical data support that genital tract levels of precursor and active forms are fairly correlated, and that vaginal PK profiles of TDF/TFV and FTC can be used to assess the potential for protection against HIV-1 transmission [15, 57]. Thirdly, the impact of sexual intercourse on PK was not considered despite being likely considerable. Fourth, the role of liposomes and their association with fibers in enhancing mucosal drug levels was not fully assessed in this work. However, our previous work pinpoints that the association of nanocarriers to polymeric platforms used for vaginal administration, namely rapid-dissolving films, potentiates the local drug retention and distribution [30]. In a related study, Krogstad *et al*. [58] reported that the incorporation of etravirine-loaded PLGA nanoparticles into PVA fibers provided enhanced vaginal retention of both nanocarrier and the antiretroviral drug over 7 days. Although the outcome of using liposomes-in-PVA fibers was not so remarkable, higher drug levels in vaginal lavage as compared to PCL fibers could be attributed to a similar effect resulting from the administration of a nanocarrier-in-fiber composite. Finally, formal safety studies were not conducted. Still, the reduced cytotoxicity of fibers, alongside the favorable safety track record of individual materials, seem to backup that PCL fibers and liposomes-in-PVA fibers are potentially safe.

## 4. Conclusions

We produced and characterized two formulations based on polymeric fibers that could be useful for the vaginal delivery of TDF and FTC in the context of topical PrEP. PCL fibers and liposomes-in-PVA fibers were shown to possess adequate biopharmaceutical properties, namely drug release profiles compatible with on-demand microbicide use and low toxicity to cells of the female genital tract. Both types of fibers provided rapid onset of local drug levels upon single vaginal administration to mice, namely when compared to the continuous use of oral TDF/FTC. Moreover, drug concentrations in vaginal fluids were moderately sustained up to 1-4 h, which could be translatable into a fairly wide protection time window in humans. This was particularly noticeable for liposomes-in-PVA fibers. The ability of fibers to provide vaginal tissue drug concentrations that were at least as high as those obtained with oral TDF/FTC was not clear and needs further clarification. Overall, the proposed drug-loaded fibers may provide an interesting alternative or a complementary tool to oral PrEP.

## Data availability

The data that support the findings of this study are available from the corresponding authors on reasonable request.

## Acknowledgements

This work was financed by Programa Gilead GÉNESE (refs. PGG/046/2015) and Portuguese funds through FCT – Fundação para a Ciência e a Tecnologia/Ministério da Ciência, Tecnologia e Ensino Superior in the framework of the project “Institute for Research and Innovation in Health Sciences” UID/BIM/04293/2019. The work was also financed by FCT in the framework of the Strategic Funding UID/FIS/04650/2019 and in the ambit of the project POCI-01-0145-FEDER-032651 and PTDC/NAN-MAT/326512017, co-financed by the FEDER, through COMPETE 2020, under PORTUGAL 2020, and FCT. Marlene Lúcio thanks FCT and ERDF for doctoral position Ref. CTTI-150/18-CF(1) in the ambit of the project CONCERT (POCI-01-0145-FEDER-032651 and PTDC/NAN-MAT/326512017).

## Author contributions

M.L., T.V., B.S. and J.d.N. conceived the overall study design and methodological approach; M.L. and J.d.N. were responsible for project management; R.N., S.B., M.J.F., H.G., M.L. and J.d.N. conducted experimental work; all authors analyzed data; M.L. and J.d.N. wrote the initial draft manuscript; all authors contributed to manuscript revision and editing; and M.L., T.V., B.S. and J.d.N. contributed to funding acquisition.

