## Supporting Information for "Electrospun fibers for vaginal administration of tenofovir disoproxil fumarate and emtricitabine in the context of topical pre-exposure prophylaxis"

### S1. Supporting Methods

#### *Biological sample processing and drug assay by liquid chromatography-tandem mass spectrometry*

**Chemicals.** TDF (MW=635.5 g·mol$^{-1}$), TFV (MW=287.2 g·mol$^{-1}$) and FTC (MW=247.3 g·mol$^{-1}$) were obtained from the suppliers indicated in the main manuscript. Isotopically-labeled compounds $^{2}H_{12}$ TDF-iso (MW=647.59 g·mol$^{-1}$) and $^{2}H_{3}$, $^{15}N$ FTC-iso (MW=251.26 g·mol$^{-1}$) were purchased from Alsachim (Illkirch-Graffenstaden, France), and $^{2}H_{6}$ TFV-iso (293.25 g.mol-1) from LGC Standards (Barcelona, Spain). LC-MS grade acetonitrile and acetic acid were acquired from Merck (Darmstadt, Germany). Ultrapure water was obtained in-house using a Milli-Q purification system (Merck Millipore, Darmstadt, Germany).

**Preparation of samples.** Samples were rapidly thawed in a water bath at 37 °C. Vaginal tissues were mixed with acetonitrile (1:10, *w/v*) and homogenized using an Ultra-Turrax processor (IKA, Staufen, Germany). In the case of vaginal lavage and blood plasma, samples were mixed with an equal volume of acetonitrile (1:1, *v/v*) and homogenized by vortexing. All samples were then centrifuged (13,414 ×*g*, 10 min, 4 °C), and supernatants were collected and filtered by 0.20 µm Millex-LG PTFE filters (Merck Millipore, Tullagreen, Ireland) before being assayed by LC-MS/MS.

**Preparation of standard solutions and mobile phase.** Stock solutions of TDF, TFV, FTC and isotopically-labeled compounds (used as internal standards) were prepared in ultrapure water at 1 mg·mL$^{-1}$ and stored at −20 °C. Intermediate solutions of each compound were prepared daily at 10,000 ng·mL$^{-1}$ and 1,000 ng·mL$^{-1}$ in ultrapure water and further diluted in order to achieve final concentrations of 0.5, 1, 5, 10, 50, 100, 200, 350, 500 and 750 ng·mL$^{-1}$ in a mixture of acetonitrile and water (5:95, *v/v*). Internal standards were added to the corresponding parent compound standard solution to a final concentration of 60 ng·mL$^{-1}$. The mobile phase comprised 0.1% (*v/v*) acetic acid in water and 0.1% (*v/v*) acetic acid in acetonitrile as aqueous and organic components, respectively. Both solutions were filtered either through a 0.22 µm Millipore GVWP filter (aqueous component) or a 0.45 µm Millipore HVHP filter (organic component), and degassed in an ultrasonic bath for 15 min prior to use.

**Chromatographic conditions.** Sample separation was conducted using an Accela HPLC (Thermo Fisher Scientific, Bremen, Germany) and a reversed phase Mediterranea Microbore C18 column (3 µm, 100 mm x 2.1 mm; Teknokroma, Barcelona, Spain) was used as stationary phase and maintained at 45 °C. The injection volume was 20 µL. Elution was performed in gradient mode at room temperature, starting with 100% of aqueous component and reaching 80% of organic component over 4.50 min at a flow rate of 0.35 mL·min$^{-1}$. The contribution of the aqueous component was changed to 100% from 4.50 to 4.60 min, and maintained for an additional 4.40 min. Retention time of 2.68 min, 3.34 min and 3.14 min for TFV, TDF and FTC, respectively. Quantitative analyses were done on an LTQ™ Orbitrap™ XL hybrid mass spectrometer (Thermo Fischer Scientific) controlled by LTQ™ Tune Plus 2.5.5 and Xcalibur™ 2.1.0 software (Thermo Fischer Scientific). The capillary voltage of the electrospray ionization source (ESI) was set to 3.1 kV. The capillary temperature was 300 °C. The sheath gas and auxiliary gas flow rates were set at 40 and 10 (arbitrary unit as provided by the software settings). The capillary voltage was 36 V and the tube lens voltage was 100 V on the positive mode. The resolution of SIM MS scan was 60,000 with center mass *m/z* of 288.05, 278.05 and 520.18, and width of 5, 5 and 10. Data dependent MS/MS was performed on CID using helium as collision gas with energy settings of 45 V. The mass spectrometer transitions (*m/z*) were 520.18>288.05 for TDF, 288.08>270.07 for TFV and 248.05>129.85 for FTC. TDF-iso, TFV-iso and FTC-iso were used as IS and monitored at *m/z* transitions 532.25>290.10, 294.12>276.06 and 252.06>132.04, respectively. MS data handling software Xcalibur™ Qual Browser (Thermo Fischer Scientific) was used to search for predicted metabolites by their *m/z* value and MS/MS value. Peak area ratio (drug/IS) versus nominal concentration of TFV, TDF and FTC were plotted to generate calibration curves using linear regression. Linearity was established over the range of 0.5-750 ng.mL$^{-1}$ for all drugs.

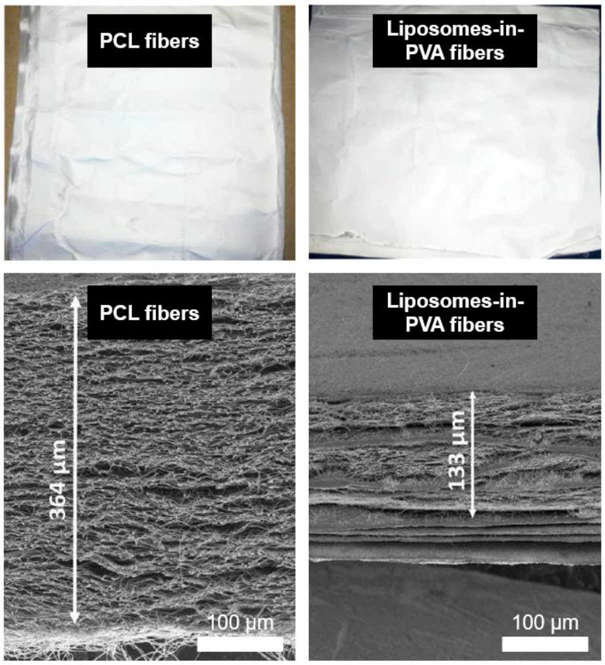


**Figure S1.** Macroscopic appearance (top images) and lateral SEM view (bottom images) of PCL fibers and liposomes-in-PVA fibers.

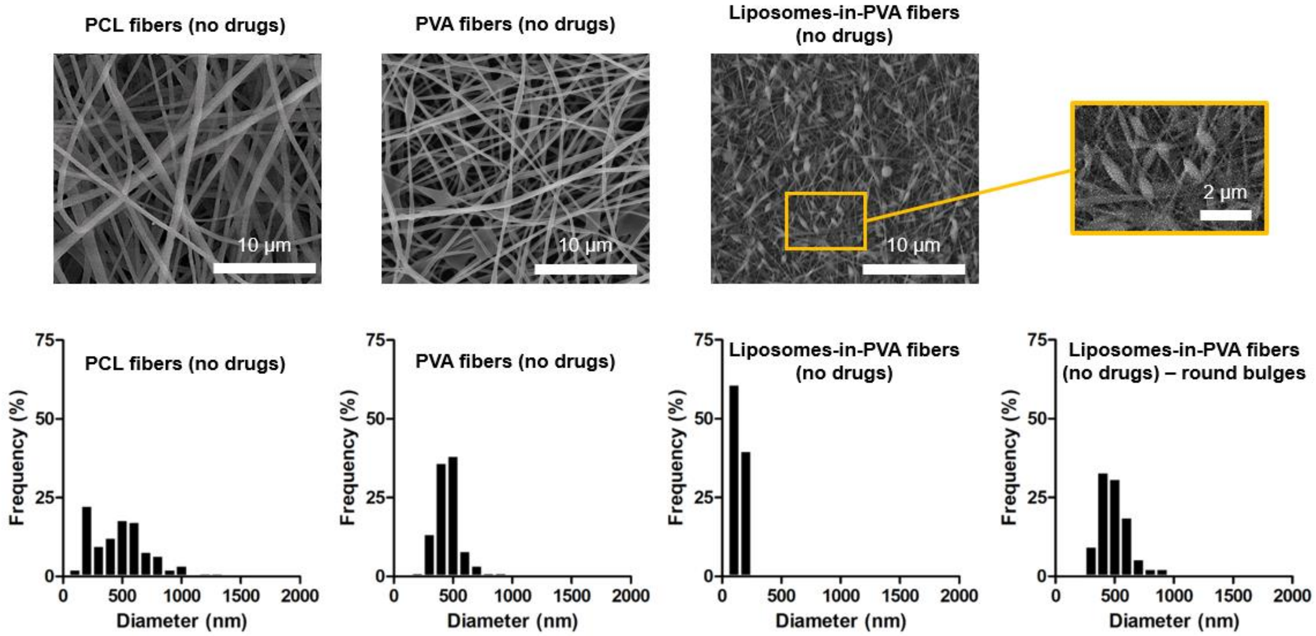


**Figure S2.** SEM images and cross-section diameter distribution of PCL fibers (no drugs), PVA fibers (no drugs) and liposomes-in-PVA fibers (no drugs). Specific data are also presented for enlarged sections (round bulges) of liposomes-in-PVA fibers (no drugs).

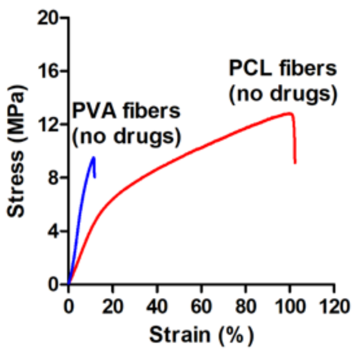


**Figure S3.** Mechanical properties of PCL fibers (no drugs) and PVA fibers (no drugs).

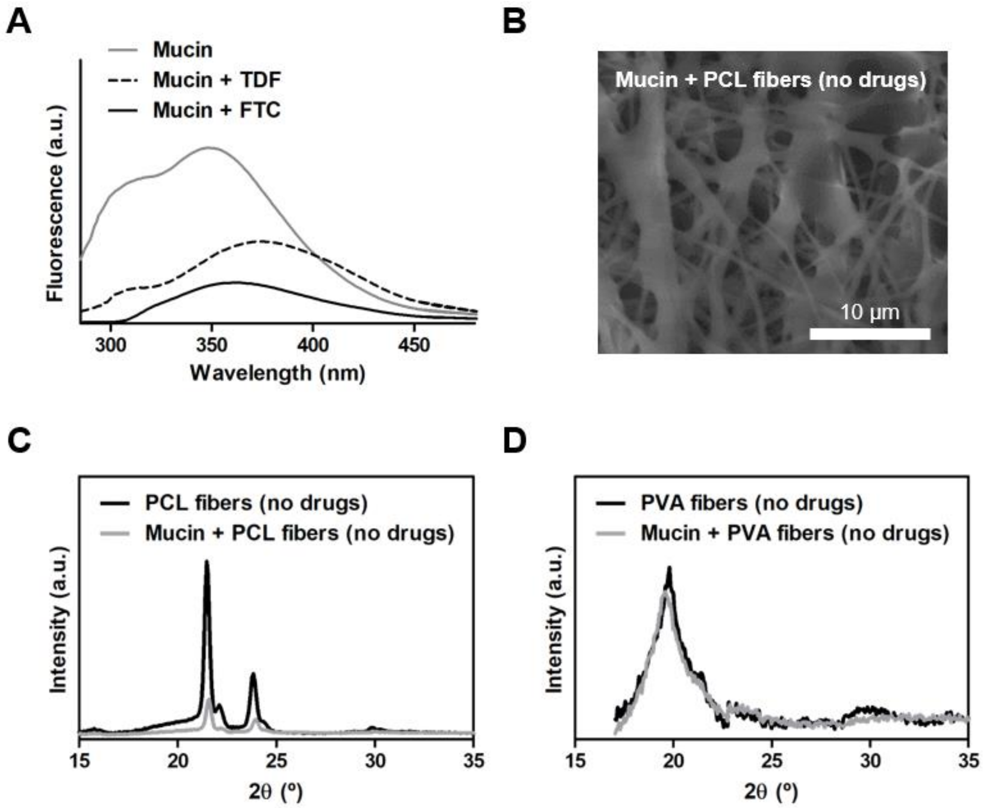


**Figure S4.** Interactions between fibers and mucin. **(A)** Mucin fluorescence quenching by TDF and FTC. **(B)** SEM image of PCL fibers (no drugs) after incubation with mucin. X-ray diffractograms of **(C)** PCL fibers (no drugs) and **(D)** PVA fibers (no drugs) in the presence or absence of mucin.

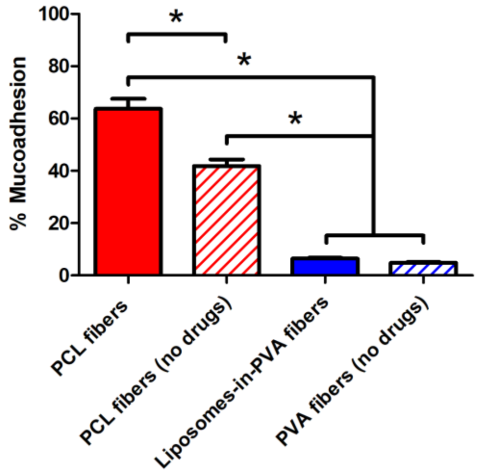


**Figure S5.** Estimation of the mucoadhesive potential of different fibers. Results are presented as mean ± SD (*n* = 3). (∗) denotes statistically significant differences ($p < 0.05$).

**Table S1.** Constant rates (*k*) and adjusted correlation coefficients ($R^2_{adjusted}$) obtained by fitting FTC and TDF release profiles from PCL fibers to different kinetics models.

| Drugs | First-order model | | Gallagher-Corrigan model | | Korsmeyer-Peppas model | | |
|---|---|---|---|---|---|---|---|
| | $k$ | $R^2_{adjusted}$ | $k$ | $R^2_{adjusted}$ | $k$ | $n$ [(a)] | $R^2_{adjusted}$ |
| TDF | 33.2 ± 10.9 | 0.293 | 24.0 ± 2.1 | 0.981 | 51.9 ± 1.6 | < 0.5 | 0.884 |
| FTC | 36.8 ± 12.7 | 0.287 | 30.9 ± 1.1 | 0.982 | 81.5 ± 3.5 | < 0.5 | 0.936 |

[(a)] Release exponent parameter (*n*) below 0.5 is indicative of Fickian diffusion.

**Table S2.** Constant rates (*k*) and adjusted correlation coefficients ($R^2_{adjusted}$) obtained by fitting FTC and TDF release profiles from liposomes-in-PVA fibers to different kinetics models.

| Drugs | First-order model | | Gallagher-Corrigan model | | Korsmeyer-Peppas model | | |
|---|---|---|---|---|---|---|---|
| | $k$ | $R^2_{adjusted}$ | $k$ | $R^2_{adjusted}$ | $k$ | $n$ [(a)] | $R^2_{adjusted}$ |
| TDF | 9.8 ± 1.0 | 0.976 | 9.8 ± 1.3 | 0.979 | 89.2 ± 2.1 | < 0.5 | 0.891 |
| FTC | 7.1 ± 1.3 | 0.986 | 6.5 ± 1.1 | 0.989 | 70.8 ± 2.7 | < 0.5 | 0.954 |

[(a)] Release exponent parameter (*n*) below 0.5 is indicative of Fickian diffusion.

**Table S3.** Number of mice (out of 5) presenting readily identifiable residues of fiber mats in the vagina during necropsy.

| Time points | PCL fibers | Liposomes-in-PVA fibers |
|---|---|---|
| 15 min | 4 | 2 |
| 1 h | 3 | 1 |
| 4 h | 1 | 0 |
| 24 h | 0 | 0 |
